# Genome-wide variation in DNA methylation linked to developmental stage and chromosomal suppression of recombination in white-throated sparrows

**DOI:** 10.1101/2020.07.24.220582

**Authors:** Dan Sun, Thomas S. Layman, Hyeonsoo Jeong, Paramita Chatterjee, Kathleen Grogan, Jennifer Merritt, Donna L. Maney, Soojin V. Yi

**Affiliations:** School of Biological Sciences, Institute for Bioengineering and Bioscience, Georgia Institute of Technology, Atlanta, GA 30332; Department of Epigenetics and Molecular Carcinogenesis, The University of Texas MD Anderson Cancer Center, Smithville, TX 78957; Department of Psychology, Emory University, Atlanta, GA 30322; Departments of Anthropology & Biology, University of Cincinnati, Cincinnati, OH 45221

**Keywords:** DNA methylation, recombination, gene expression, epigenetic reprogramming, transposable elements

## Abstract

DNA methylation is known to play critical roles in key biological processes. Most of our knowledge on regulatory impacts of DNA methylation has come from laboratory-bred model organisms, which may not exhibit the full extent of variation found in wild populations. Here, we investigated naturally-occurring variation in DNA methylation in a wild avian species, the white-throated sparrow (*Zonotrichia albicollis*). This species offers exceptional opportunities for studying the link between genetic differentiation and phenotypic traits because of a non-recombining chromosome pair linked to both plumage and behavioral phenotypes. Using novel single-nucleotide resolution methylation maps and gene expression data, we show that DNA methylation and the expression of DNA methyltransferases are significantly higher in adults than in nestlings. Genes for which DNA methylation varied between nestlings and adults were implicated in development and cell differentiation and were located throughout the genome. In contrast, differential methylation between plumage morphs was localized to the non-recombining chromosome pair. One subset of CpGs on the non-recombining chromosome was extremely hypomethylated and localized to transposable elements. Changes in methylation predicted changes in gene expression for both chromosomes. In summary, we demonstrate changes in genome-wide DNA methylation that are associated with development and with specific functional categories of genes in white-throated sparrows. Moreover, we observe substantial DNA methylation reprogramming associated with the suppression of recombination, with implications for genome integrity and gene expression divergence. These results offer an unprecedented view of ongoing epigenetic reprogramming in a wild population.

## INTRODUCTION

DNA methylation is a key epigenetic mark that adds another layer of information to the genomic DNA (1). The best-known effect of DNA methylation, which has been observed most often in mammalian studies, is the repression of transcription resulting from methylation at CpG sites in *cis*-regulatory regions (2). The role of DNA methylation in other genomic regions is less well understood, although there are examples linking DNA methylation of gene bodies and intergenic regions to regulation of gene expression (3-5). DNA methylation in non-genic regions has been implicated in many regulatory processes, including maintenance of genome stability and silencing transposable elements (TEs) (6-9). DNA methylation is also known to associate with development and aging (10-14).

Most of our knowledge about DNA methylation comes from studies of humans and laboratory mice. Little is currently known about how DNA methylation varies and how it impacts gene expression in natural populations. In this study, we provide rare insight into how DNA methylation varies in a wild species of songbird. We used deep whole-genome bisulfite sequencing (WGBS) to generate single-nucleotide-resolution maps of DNA methylation in a wild passerine, the white-throated sparrow (*Zonotrichia albicollis*). This species is an exceptional non-model vertebrate system for understanding links between the evolution of genomes and complex behavioral phenotypes (15-17). Two naturally occurring plumage morphs, white-striped (WS) and tan-striped (TS), are completely linked to a supergene that segregates with an aggressive phenotype in both sexes. Birds of the WS morph, which are heterozygous for a rearranged second chromosome (ZAL2^m^), are on average more aggressive than birds of the TS morph (18,19), which are homozygous for the standard arrangement (ZAL2) (20-22). In addition, WS birds invest less in parenting behavior than do TS birds (18,19,23-27). Thus, the supergene is associated with a complex phenotype involving both aggression and parenting.

This unique chromosomal polymorphism is maintained in the population through disassortative mating; that is, nearly all mating pairs consist of one TS and one WS individual (15,21,22). As a consequence, the ZAL2^m^ chromosome is nearly always in a state of heterozygosity, experiencing little recombination. Cessation of recombination causes several genetic changes, including reduction of gene expression, accumulation of transposable elements, and ultimately, genetic degeneration of the non-recombining region (28,29). The ZAL2 and ZAL2^m^ chromosomes are in an early stage of genetic differentiation, having accumulated approximately 1% nucleotide divergence (15,16,20). The ZAL2^m^ chromosome has yet to exhibit signs of substantial genetic degeneration (i.e., only a handful of genes have become pseudogenized, (16)). Despite this modest genetic divergence, genes on the non-recombining ZAL2^m^ chromosome exhibit reduced expression, and ZAL2 appears to have evolved incipient dosage compensation, indicating rapid regulatory evolution preceding large-scale genetic differences between the ZAL2 and ZAL2^m^ chromosomes (16). Our novel whole genome DNA methylation maps offer a unique opportunity to investigate how DNA methylation changes in the early stage of chromosomal differentiation following the cessation of recombination.

We investigated patterns of DNA methylation in 12 samples of brain tissue from female white-throated sparrows of both morphs. These samples were taken from seven adults (four WS and three TS) and five nestlings (three WS and two TS), thus spanning two developmental time points. Our novel and comprehensive data on nucleotide-resolution whole-genome methylation maps reveal previously unknown epigenetic variation in a wild avian species. We find substantial variation in DNA methylation between developmental stages and plumage morphs. By integrating this dataset with novel gene expression data taken from the same individuals, as well as an open chromatin map of a WS bird, we demonstrate that variation in DNA methylation between nestlings and adults is widespread across the genome and correlated with variation in expression of developmental genes. Furthermore, by identifying allele-specific methylation and its potential evolutionary origins, we provide a rare glimpse into epigenetic reprogramming of a chromosome following a recent cessation of recombination.

## MATERIALS AND METHODS

### Sample collection

For WGBS and RNA-seq experiments, we collected 12 female birds (seven breeding adults and five nestlings) for our analysis (Supplementary Table 1). Adults were collected using mist nets at our field site near Argyle, Maine, USA, as previously described (18,27). Nestlings were collected from nests at day 7 post-hatch (30). Only one nestling per nest was used for the analysis. The hypothalamus was micro-dissected from each brain as previously described (27). For the ATAC-seq experiment, the hypothalamus was micro-dissected from a non-breeding WS male adult bird (sample ID: ID 17031) collected at our field site in Atlanta, Georgia, USA (31). We performed the kinship analysis using KING (32). The kinship coefficients between the 12 individuals in this study were all practically zero (the maximum kinship coefficient was 0.00277), indicating that they were not closely related.

### Whole genome sequencing

Whole genome sequencing libraries were generated from DNA extracted from the white-throated sparrow livers using a QIAGEN DNeasy Blood and Tissue DNA kit. For each sample, 500ng-1µg of DNA was extracted and sheared on a Covaris ultrasonicator to 200-600bp at the Emory Integrated Genomics Core. The DNA fragment ends were repaired, and A-overhangs were added before Nextera barcode adaptors were ligated to the DNA fragments overnight. Finally, the libraries were PCR-amplified to increase concentration and enrich for adaptor-ligated DNA fragments. WGS libraries were sequenced using Illumina HiSeq X Ten with 150 × 2 bp paired-end reads at Macrogen Clinical Laboratory.

### SNP calling and identification of fixed differences

To identify SNPs occupying CpG sites, we first removed adaptor sequences and low-quality bases from the sequencing reads using the parameters “-q 30 -O 1 -m 50 --trim-n --pair-filter any” using cutadapt 1.18 (33). Trimmed reads were then aligned to the TS reference genome using Bowtie2 v2.3.4.2 (34) with the --very-sensitive-local option, and the alignment rate was ∼95% per sample. Technical duplicates were then discarded by Picard Tools 2.19.0 (https://broadinstitute.github.io/picard/). SNP calling was conducted on clean and aligned reads using GATK 4.0 (35-37). Specifically, SNPs were called using Haplotypecaller with the -ERC GVCF option, and joint genotyping of all samples was performed with the GenotypeGVCF. Finally, SNPs with MAF < 0.05, meanDP < 5 and meanDP > 80 were discarded using VCFtools 0.1.15 (38).

With the final set of SNPs, we identified putatively fixed differences between ZAL2 and ZAL2^m^ using the same procedure as described by Sun et al. (2018). For further alignment of WGBS, ATAC-seq, and RNA-seq data, to minimize potential mapping bias towards the reference genome (ZAL2/ZAL2) caused by differences between ZAL2 and ZAL2^m^, we constructed a genome with putatively fixed differences masked by *N*’s in the reference (*N*-masked genome).

### Whole genome bisulfite sequencing

WGBS libraries were prepared using a custom protocol. First, DNA was extracted from the hypothalamus samples using a QIAGEN DNeasy Blood and Tissue DNA kit. For each sample, 100 ng - 1 µg of DNA was pooled with 1-5% lambda phage DNA to test for bisulfite conversion efficiency. The DNA samples were then sheared on a Covaris ultrasonicator to 200-600bp. The DNA fragment ends were repaired, and A-overhangs were added before bisulfite compatible adaptors were ligated to the DNA fragments overnight. Then, the DNA fragments were bisulfite-converted and PCR-amplified to increase concentration and enrich for adaptor-ligated DNA fragments. WGBS libraries were then sequenced using Illumina HiSeq X Ten at Macrogen Clinical Laboratory. At least ∼100 million 150 bp x 2 raw reads were generated per sample (Supplemental Table 1).

### Analysis of whole genome bisulfite sequencing data

WGBS reads were trimmed as described above. The trimmed reads were aligned to the *N*-masked reference genome with parameters “--bowtie2 -X 1000” using Bismark v0.20.0 (39). The average mapping efficiency of samples was ∼70% for all samples (Supplemental Table 1). Third, duplicated reads and non-bisulfite-converted reads were discarded by *deduplicate_bismark* (parameter: -p) and *filter_non_conversion* (parameter: percentage_cutoff 20), respectively. Last, *bismark_methylation_extractor* was run to extract CpG methylation calls. To obtain bisulfite conversion rates, raw reads were aligned to the phage lambda genome using Bismark (same parameters). Because lambda DNA is not methylated and therefore should be completely bisulfite-converted, the percentage of methylated cytosines of lambda DNA was taken as the non-conversion rate. Bisulfite conversion rates were above 99.8% in all samples (Supplemental Table 1).

To call allele-specific methylation values, SNPsplit 0.3.4 (40) was run with parameters “--bisulfite --paired” using fixed differences between ZAL2 and ZAL2^m^. Then, *bismark_methylation_extractor* was run for allele-separated reads. For WS birds, consistent with the genotype (ZAL2/ZAL2^m^), the percentage of reads assigned to each chromosome was ∼4 - 4.5% (Supplemental Table 1); for TS birds, the percentage of reads assigned to ZAL2 was ∼8 - 9% but to ZAL2^m^ 0 - 0.01% (Supplemental Table 1), which was consistent with the genotype (ZAL2 / ZAL2). After this procedure, the median sequencing depths were at least 9 reads per sample and 4 per allele (Supplemental Table 1). Only CpG sites with at least five reads aligned were retained for further analysis (e.g., (41)). Finally, because cytosine polymorphisms could hamper accurate calling of methylation, we excluded any CpGs in the reference genome that were polymorphic within the sequenced samples.

### ATAC-seq library preparation, sequencing, data pre-processing, and peak calling

For one sample (hypothalamus of a WS male, ID 17031), 10,000 - 200,000 cells were homogenized in EMEM (Eagle’s Minimum Essential Medium) and phosphate-buffered saline. The cells were pelleted in a centrifuge and re-suspended in a lysis buffer made of non-ionic detergent (made in-house from Tris, NaCl, MgCl_2_, and IGEPAL CA-630). After cell lysis, nuclei were isolated by centrifugation and added to a tagmentation reaction mix (Illumina Nextera DNA Library Prep Kit, Cat#: FC-121-1030). During tagmentation, the sequencing adapters were inserted into accessible chromatin regions by Tn5 transposase. Adapter-tagmented fragments were purified (Invitrogen Agencourt AMPure XP beads, Cat#: A63880), bar-coded (Illumina Nextera Index Kit, cat#: FC-121-1011), and amplified (Fisher KAPA HiFi HotStart Kit, Cat#: NC0295239). The ATAC-seq libraries were then sequenced using a MiSeq sequencer (Illumina; Reagent Kit v3) with 150 cycles (75 bp paired-end reads) in the Molecular Evolution Core at Georgia Tech.

We aligned the trimmed ATAC-seq reads (trimming was performed as above) to the *N*-masked reference genome using Bowtie2 v2.3.4.2 (parameters: -X 2000 --no-mixed --no-discordant) (34), which allowed a maximal insert size of 2 Kb between paired reads, and discarded unmapped or discordant alignments. The mapping efficiency for this sample was 82.13%. The aligned reads were then deduplicated using markdup of SAMtools 1.7 (42). As a result, we obtained 23 million clean mapped reads. To identify ZAL2 and ZAL2^m^-specific ATAC-seq peaks, we followed the strategy proposed in (43). Specifically, we first called peaks in the overall sample using MACS2 version 2.1.1.20160309 (44) with ‘-g 1.1e+9 -f BAMPE -p 0.05 -B --SPMR --nomodel’ options. We next assigned reads to ZAL2 and ZAL2^m^ using SNPsplit 0.3.4 (40) with parameters “--paired” using fixed differences between ZAL2 and ZAL2^m^. The number of ZAL2 and ZAL2^m^ reads mapped to the ATAC-seq peaks were counted using Bedtools v2.28.0 (45), and the differences in allelic read counts were tested by a two-tailed binomial test. Peaks with FDR-corrected *P* < 0.05 were denoted as allele-specific.

### RNA-seq library preparation, sequencing, data processing, and analysis of differential expression

RNA extraction and library preparation of the female samples were performed as previously described (27). The libraries were then sequenced on the HiSeq 4000 at 150 bp paired-end reads to ∼40 million reads per sample. RNA-seq raw reads were trimmed as above and then aligned to the *N*-masked genome by HISAT2 2.1.0 (46). Secondary alignments were filtered by SAMtools 1.7 (42) to ensure that only primary alignments were retained. SNPsplit 0.3.4 (40) was run to assign reads to ZAL2 or ZAL2^m^ for the WS samples. Expression levels (raw read counts) were then quantified by StringTie v1.3.4d (47). To identify genes that were differentially expressed between ZAL2 and ZAL2^m^, we normalized libraries with the size factors generated in the morph comparison step and identified differential expression with ‘design = ∼ age + allele’ (age as the adjusted covariate) using the DESeq2 1.22.2 package (48) in R 3.5 (49).

### Analysis of differential DNA methylation

Differentially methylated CpGs between age groups and morphs (or alleles) were detected by DSS 2.30.1 (50) under the default setting, with one variable as the ‘independent’ variable and the other as ‘adjusted’ covariate. CpGs with FDR-corrected *P*-values less than 0.05 and absolute values of differences in methylation greater than 10% were defined as DMCs. Bedtools v2.28.0 (45) was run to assign DMCs to different gene features. If a DMC was within multiple gene features, we prioritized the assignment in the following order: upstream (10 Kb upstream of TSSs), exons, introns, downstream (10 Kb downstream of TESs), TEs and intergenic regions. After quality control, a total of 3,880,473 CpGs were used to identify age-DMCs and morph-DMCs, and 317,499 CpGs were used to identify allele-DMCs.

### Principal component analysis

We stored DNA methylation data generated from all samples as a methylrawDB object using methylkit 1.9.4 (51). The object was then converted into a percent methylation matrix, with only CpG sites with more than five reads in all samples retained. PCA analysis was performed using the PCASamples function in methylkit (parameter: obj.return = T). The returned prcomp result was used to plot sample clusters with the autoplot function in ggfortify 0.4.5 (52).

### Transposable element annotation

We adopted both *de novo* and homology-based approaches to annotate repetitive sequences in the reference genome. First, *de novo* discovery of TEs was performed by RepeatModeler 1.0.9 (53). The generated library was merged with the avian Repbase library (20181026 version), which was used to annotate TEs in the reference genome using RepeatMasker 4.0.9 (parameters: -xsmall -s -nolow -norna -nocut) (54).

### Cross-species whole genome alignment and comparison of DNA methylation

We examined cross-species alignment to identify CpGs specific to ZAL2 and ZAL2^m^. We aligned the sparrow reference genome to the zebra finch (Taeniopygia_guttata-3.2.4) and great tit (Parus_major1.1) reference genomes using minimap2-2.16 (parameters: -- secondary=no -c) (55). Only alignments for which we were confident, defined by the highest mapping score (MAPQ=60), were retained. The paftools liftover program of minimap2 was then run to find dinucleotides in the great tit genome that were orthologous to CpG sites in the sparrow genome. Note that chromosome number in white-throated sparrows follows the conventional nomenclature for avian chromosomes, numbering them from largest to smallest (56). Chromosome 2 in white-throated sparrows corresponds to chromosome 3 in chicken (20). The inter-species alignments were consistent with the homology between Chromosome 3 in zebrafinch-ZAL2 in white-throated sparrows. In addition, Using the brain methylation data from the great tit compiled by Sun et al. (57), we obtained fractional methylation levels for shared CpGs in the other two species (sparrow-tit: 436 CpGs).

## RESULTS

### Contrasting effects of developmental stage and morphs on genome-wide DNA methylation maps

We examined patterns of DNA methylation and gene expression in samples of hypothalamus, a brain region known to take part in many social and developmental traits. Our experimental cohort included birds of both morphs in two age groups: adults aged more than one year (7 WS, 5 TS) and nestlings at post-hatch day seven (3 WS, 2 TS). To remove the effect of sex in a cost-effective manner, in the current study only female birds are analysed for whole genome DNA methylation. We investigated the effects of morph and developmental stage (also referred to as ‘age’ in this manuscript) on genome-wide DNA methylation maps. These white-throated sparrows were not related based on the kinship coefficient analysis (32). Specifically, using all SNPs we detected in this study, the maximum kinship coefficient was 0.00277 (Materials and Methods).

Whole genome bisulfite sequencing maps of the samples were generated (see Methods). Bisulfite conversion rates, determined from spiked-in unmethylated lambda phage DNA, were all 99.8%. We generated on average 431 million reads of 150 bps for WS birds, and 134 million reads for TS birds. Greater coverage for the heterozygous WS birds was necessary to recover sufficient reads for both ZAL2 and ZAL2^m^ chromosomes. After removing duplicates, we mapped the reads to an *N*-masked reference genome to avoid mapping bias due to the polymorphisms between ZAL2/ZAL2^m^ chromosomes. Following these procedures, WS birds and TS birds had on average 33.4X and 13X coverage, respectively, per CpG (Supplementary Table 1). We aimed for higher depth for the WS birds so that we can separate ZAL2 and ZAL2^m^ chromosomes for later analyses.

To first gain an understanding of genome-wide variation in DNA methylation, we performed a principal component analysis (PCA) of all mapped CpGs prior to separating the ZAL2 and ZAL2^m^ alleles (**Figure 1A**). The first principal component (PC1), which distinguished adults from nestlings, accounted for the largest amount of variation in DNA methylation (∼20%). The second principal component (PC2) separated the TS and WS morphs and explained ∼11% of the variation among samples. We then performed the same analyses using only CpGs within the rearranged portion of the ZAL2/ZAL2^m^ chromosome (**Figure 1B**), which produced highly similar results. Consequently, age and morph were determined to be the top two factors of variation in DNA methylation in our data.

**Figure 1.**
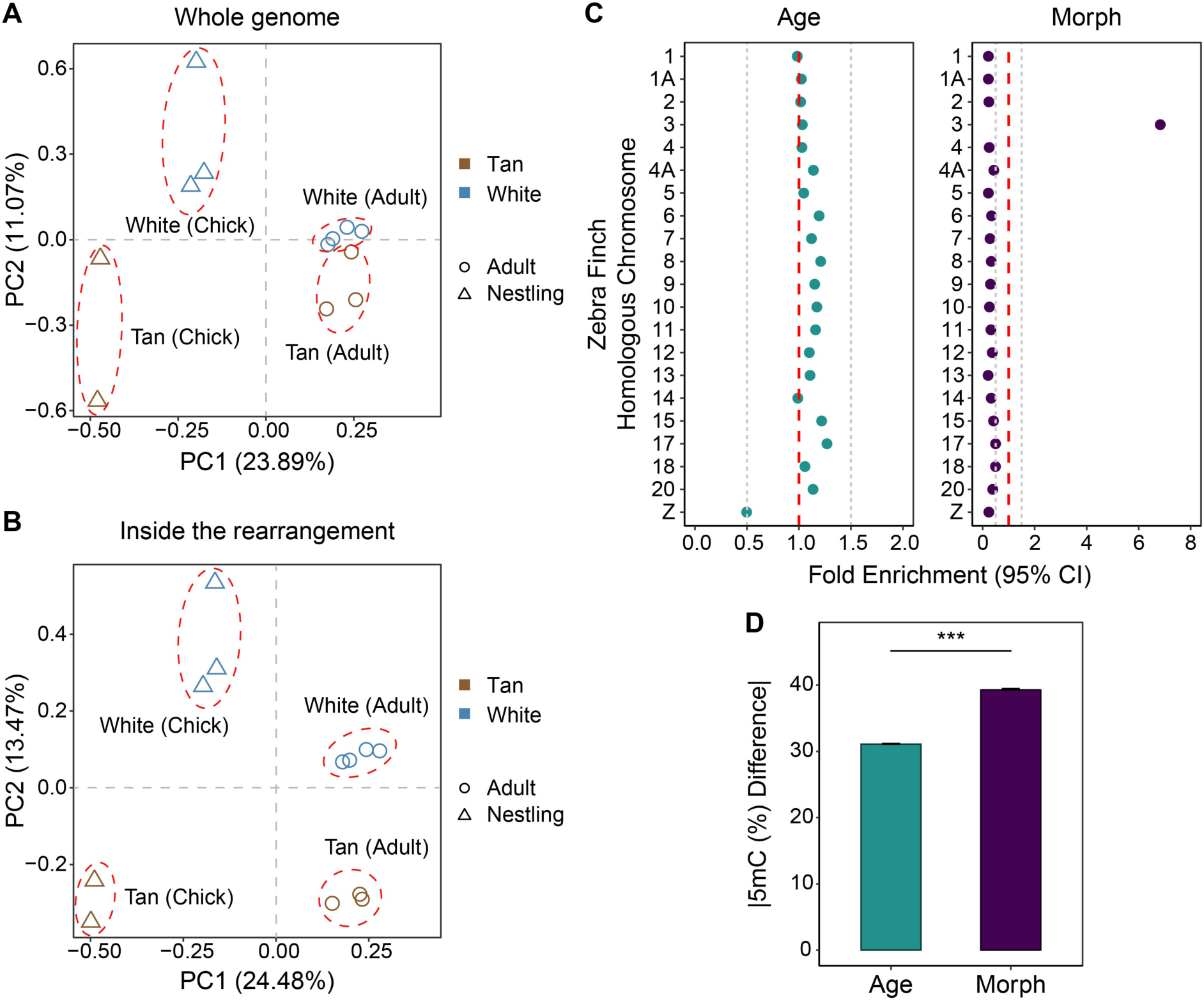
The effects of age and morph on DNA methylation patterns. PCA of WGBS samples for **(A)** all CpGs and **(B)** CpGs within the rearranged portion on ZAL2/ZAL2^m^. **(C)** Fold enrichment with 95% confidence interval (95% CI) for the chromosomal distribution of DMCs (using homologous chromosomes in zebra finch for designation). The fold enrichment and confidence intervals were calculated by comparing the real distribution of DMCs with the null distribution generated by 100 random selections of the same number of CpGs. The red dashed lines indicate no depletion/enrichment (enrichment score = 1) of DMCs on a chromosome, and the gray dashed lines depict boundaries for moderate depletion (0.5) or enrichment (1.5) of DMCs. Only chromosomes larger than 10 Mb are shown. **(D)** Mean absolute differences (with 95% confidence intervals) in fractional DNA methylation (5mC [%]) for age-DMCs and morph-DMCs. Effect sizes were smaller for age-DMCs than for morph-DMCs. \*\*\**P* < 0.001; Mann-Whitney *U* test. Standard error bars are shown.

Using the same whole-genome CpG data, we identified significantly differentially methylated CpGs (herein referred to as ‘**DMC**s’) between adults and nestlings, as well as between WS and TS birds, using a method designed specifically for the WGBS analysis (50). This method explicitly accounts for the characteristics of next-generation sequencing data and allows us to identify sites that are affected by different co-variates. In addition to correcting for multiple testing using the FDR method (FDR-adjusted *P* < 0.05), we restricted the value of the absolute methylation difference to be equal to or greater than 10%. Following these procedures, we identified 286,434 DMCs between adults and nestlings (referred to as ‘age-DMCs’), and 4,507 DMCs between TS and WS birds (referred to as ‘morph-DMCs’).

Age-DMCs and morph-DMCs were distinct from each other with respect to both the chromosomal distribution and the effect sizes (**Figure 1C** and **1D**). In terms of the chromosomal distribution, age-DMCs were distributed across the genome, but depleted from the Z chromosome. In comparison, morph-DMCs were largely restricted to the ZAL2/ZAL2^m^ chromosomes (**Figure 1C**), indicating that nearly all differences in DNA methylation between the morphs were due to CpGs on the non-recombining chromosomal pair. The effect sizes, measured as absolute differences in DNA methylation between the two morphs, were on average substantially greater than for the age-DMCs (**Figure 1D**).

### Global hypermethylation of CpGs in adults relative to nestlings

Interestingly, most age-DMCs (97.7% of all age-DMCs) were more highly methylated (hyper-methylated) in adults than in nestlings (**Figure 2A**). We examined the expression levels of DNA methyltransferases (DMNTs) in RNA-seq data of the same individuals. Consistent with the observed genome-wide hypermethylation of samples from adults, DNA methyltransferases DNMT1 and DNMT3b had significantly higher expression in adults than in nestlings (**Figure 2B**; note that DNMT3a is not annotated in the reference genome due to the poor assembly quality around that region). Genes harboring age-DMCs in promoters were significantly enriched for gene ontology (GO) terms related to development and cell differentiation (**Figure 2C**).

**Figure 2.**
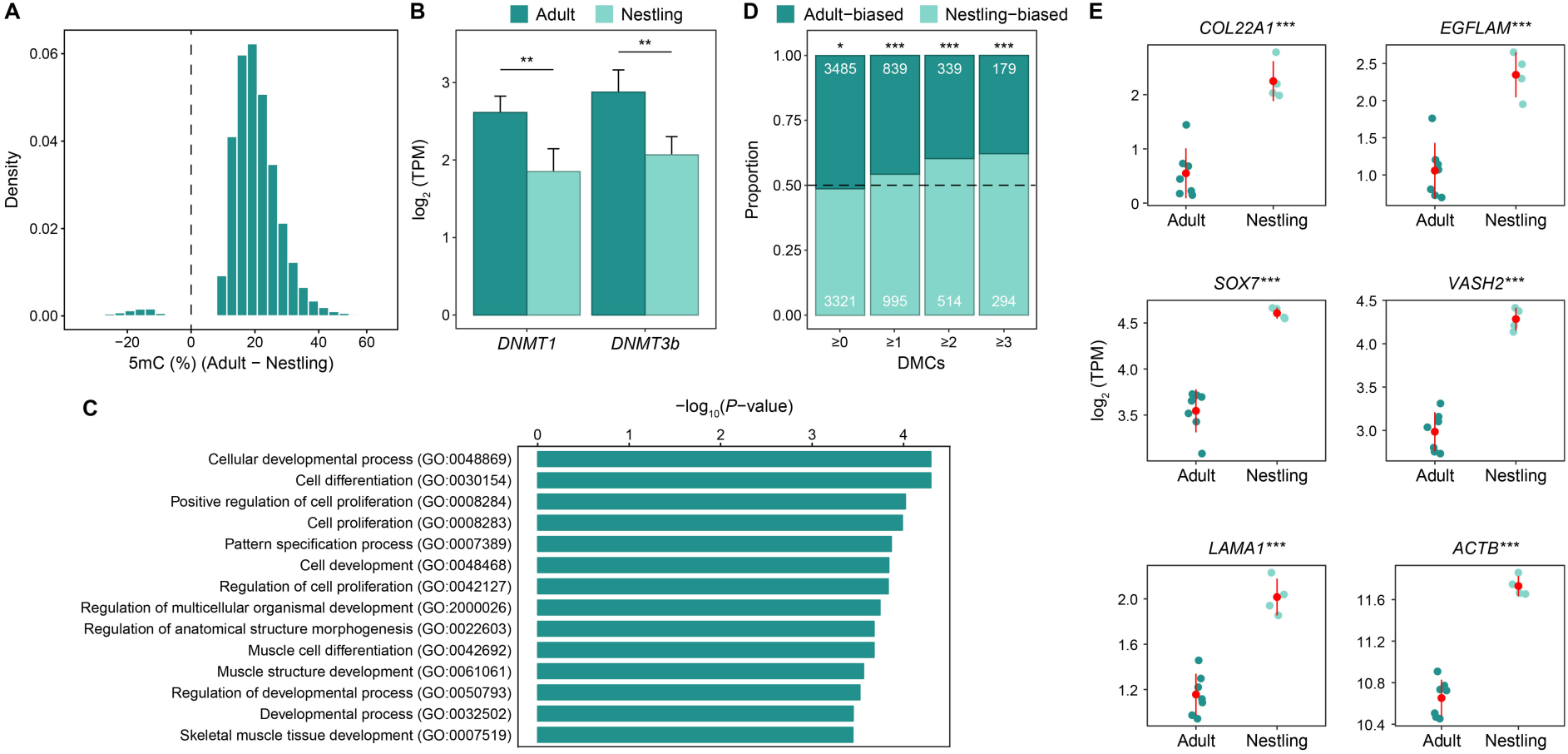
Hypermethylation in adults relative to nestlings. **(A)** The density distribution of differences in methylation between adults and nestlings shows that most age-DMCs are hypermethylated in adults, compared with nestlings. **(B)** Both *DNMT1* and *DNMT3b* were more highly expressed in adults than in nestlings (tested by DESeq2, ***:*P* < 0.001), consistent with the observed hypermethylation in adults. **(C)** The proportions of adult-biased and nestling-biased genes with more than 1, 2, or 3 age-DMCs in their promoters. The numbers of DE genes that are biased in each age group are marked. The differences in the number of DE genes between adults and nestlings were tested by a binomial test (***:*P* < 0.001). **(D)** GO enrichment of genes that contain at least three age-DMCs within their promoters (defined as within 1.5 Kb upstream of TSS). A statistical overrepresentation test was performed by PANTHER14.1 (Fisher’s exact test), with all white-throated sparrow genes present in the *Gallus gallus* annotation database as the reference list. Only GO terms with FDR-adjusted *Q* < 0.05 and fold enrichment > 1.5 are reported. **(E)** Adults in general have lower gene expression levels than nestlings for age-DE genes (tested by DESeq2, ***:*P* < 0.001) associated with developmental processes (GO:0032502) and which have at least three age-DMCs in the promoters. Shown here are some examples. Each dot represents a sample with both WGBS and RNA-seq data. Mean +/-standard deviations are depicted as red lines.

We identified a total of 6806 genes that were differentially expressed between nestlings and adults using FDR-adjusted *P* < 0.05, demonstrating that gene expression profiles change dramatically between the two developmental stages. Among these genes, a slightly greater number was more highly expressed in adults than in nestlings (3485 adult-biased versus 3321 nestling-biased). In contrast, genes harboring age-DMCs in promoters tended to be more highly expressed in nestlings, and this trend increased as the number of age-DMCs in each promoter increased (**Figure 2D**). These observations suggest that hypermethylation of promoter CpGs might contribute to the down-regulation of early developmental genes in adults. This model is further supported by the observation that several developmental genes harbored DMCs in promoters and showed reduced expression in adults compared to nestlings (**Figure 2E**).

### Differential methylation of the ZAL2 and ZAL2^m^ chromosomes is driven by substantial hypomethylation of CpGs on the non-recombining ZAL2^m^

Because the effects of morph on DNA methylation were nearly exclusive to the ZAL2/ZAL2^m^ chromosomes, we next investigated DNA methylation patterns of these two chromosomes more deeply. To do so, we used WGBS data from WS individuals and separated the ZAL2 and ZAL2^m^ alleles (see Methods). We then used the Dispersion Shrinkage for Sequencing data (DSS) package, v.2.30.1 (50) to detect CpGs that were differentially methylated between ZAL2 and ZAL2^m^ (referred to as ‘allele-DMCs’). We identified 13,773 allele-DMCs using the same criteria we used in the genome-wide analysis (FDR-adjusted *P* < 0.05, absolute methylation difference > 10%).

To examine the degree and direction of differences in DNA methylation between ZAL2 and ZAL2^m^, we first plotted the sizes of these differences (level of ZAL2^m^ methylation – ZAL2 level of methylation) in a histogram (**Figure 3A**). Their distribution revealed that allele-DMCs tended to be less methylated (also referred to as ‘hypomethylated’) on ZAL2^m^ than on ZAL2 (**Figure 3A**). We identified three distinct groups of allele-DMCs. Approximately 75% of allele-DMCs showed a difference in DNA methylation between -0.5 to 0.5, i.e. a less than 50% difference in DNA methylation between ZAL2 and ZAL2^m^ (light blue and red, **Figure 3A**). These allele-DMCs were equally likely to be more methylated on either ZAL2 or ZAL2^m^ (depicted as ‘ZAL2^m^ < ZAL2’ and ‘ZAL2^m^ > ZAL2’ in **Figure 3A**, respectively). Interestingly, the remaining 25% of allele-DMCs showed extremely differential DNA methylation, with ZAL2^m^ alleles exhibiting markedly lower DNA methylation than their ZAL2 counterparts (depicted as ‘ZAL2^m^ << ZAL2’, dark blue, in **Figure 3A**). We will refer to these three categories of allele-DMCs as ‘ZAL2^m^ hypomethylated’ (ZAL2^m^ < ZAL2, light blue in **Figure 3A**), ‘ZAL2^m^ hypermethylated’ (ZAL2^m^ > ZAL2, red in **Figure 3A**), and ‘ZAL2^m^ extremely hypomethylated’ (ZAL2^m^ << ZAL2, dark blue in **Figure 3A**) in the remainder of the paper.

**Figure 3.**
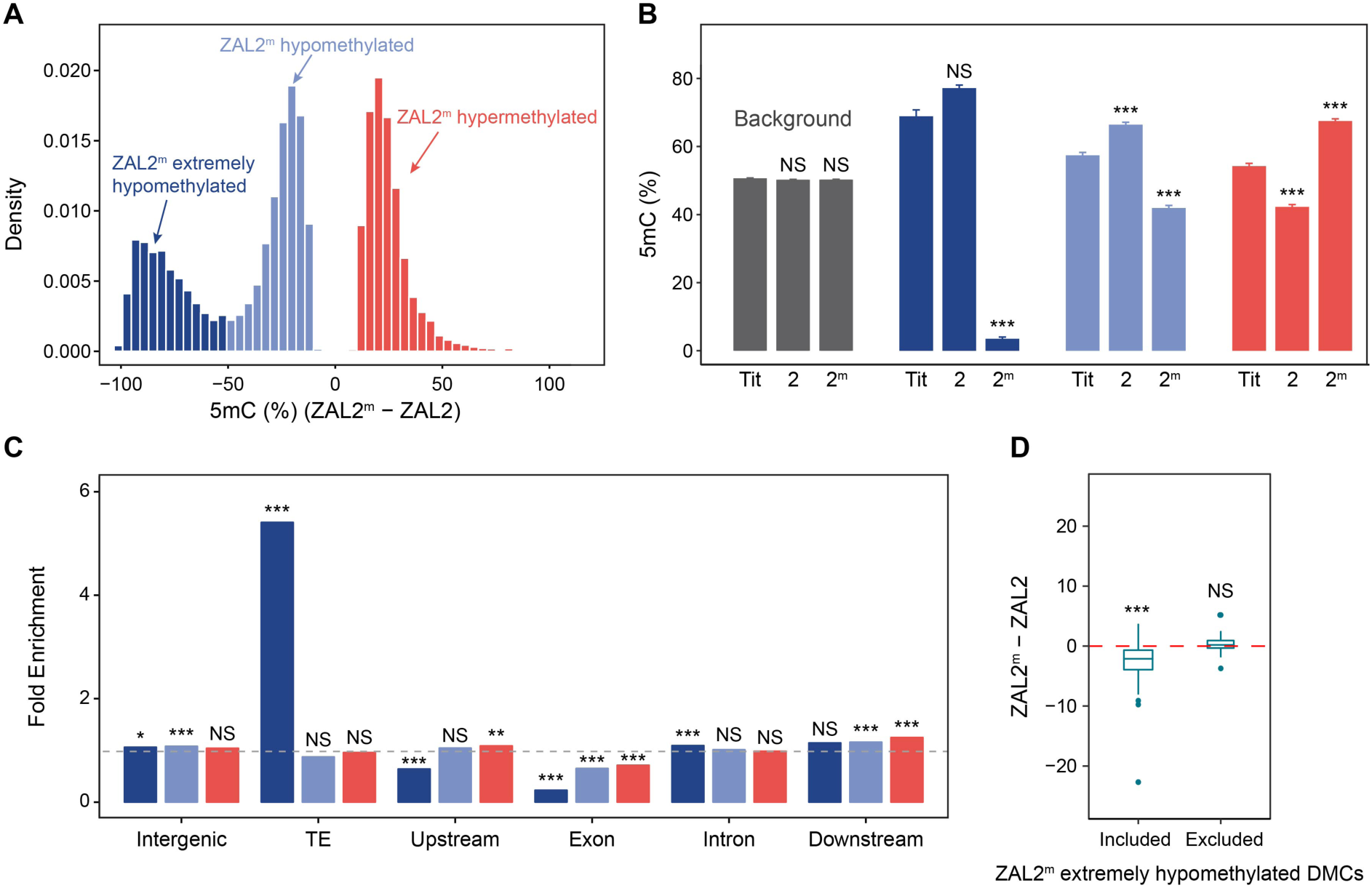
Characterization of the three classes of allele-DMCs. **(A)** The effect sizes of the differences in DNA methylation between ZAL2 and ZAL2^m^ alleles (allele-DMCs) fall into three distinct groups. **(B)** Changes in DNA methylation levels relative to the ancestral methylation levels inferred by comparison with an outgroup species, great tit. *** *P* < 0.001, Mann Whitney *U* test. **(C)** Fold enrichment of allele-DMCs within different genomic regions relative to the background (all CpGs on ZAL2/ZAL2^m^). The dashed line corresponds to a fold enrichment of 1 (no enrichment or depletion). Intergenic regions were defined as regions that were at least 10 Kb away from any genes, and upstream/downstream distal regions were defined as 10 Kb upstream/downstream of the transcription start site (TSS)/transcription end site (TES). For B-C, all ZAL2/ZAL2^m^-linked CpGs were used as the control, and enrichment or depletion was assessed by a two proportion Z-test. **(D)** Differences in TE methylation (5mC %) between ZAL2^m^ and ZAL2 after including or excluding the ZAL2^m^ extremely hypomethylated DMCs. Only TE families with more than 10 CpGs were used for analysis. For B-D, NS: not significant; \**P* < 0.05; \*\**P* < 0.01; \*\*\**P* < 0.001, Mann-Whitney *U* test.

To understand the evolutionary changes in DNA methylation leading to the three distinctive categories of allele-DMCs, we compared levels of DNA methylation in these three categories of CpGs, as well as those that did not exhibit differential DNA methylation, with corresponding levels of DNA methylation in a passerine outgroup, the great tit (58). This comparison revealed that CpGs that were not differentially methylated between the ZAL2 and ZAL2^m^ chromosomes showed similar methylation levels in the white-throated sparrow and great tit, suggesting that they have maintained similar levels of DNA methylation through evolutionary time (**Figure 3B**, gray columns). In comparison, ZAL2^m^ extremely hypomethylated (ZAL2^m^ << ZAL2) DMCs bore a clear signature of hypomethylation on the ZAL2^m^ since the split from the great tit (**Figure 3B**). This pattern contrasts clearly with that of other allele-DMCs, which exhibited signs of both increased and decreased DNA methylation compared with great tit (**Figure 3B**, light blue and red). Together these observations indicate that although both ZAL2 and ZAL2^m^ have undergone changes in DNA methylation since the divergence from the great tit, a number of CpGs on ZAL2^m^ have experienced a strong reduction in DNA methylation since the split of the ZAL2 and ZAL2^m^ chromosomes.

We then tested whether allele-DMCs are enriched in specific functional regions. While the occurrence of other allele-DMCs was similar to all CpGs, ZAL2^m^ extremely hypomethylated allele-DMCs were five-fold enriched in TEs (*P* < 2.2 × 10^−16^ using a proportion test, **Figure 3C**). They were also slightly enriched in intronic regions, while slightly (yet significantly) depleted in regions upstream of transcription start sites (TSSs) where CpG islands are typically located (e.g., (59), **Figure 3C**). Currently, TEs in white-throated sparrow are poorly annotated. We used a *de novo* annotation (see Methods) and identified subfamilies of TEs (we could not identify individual TEs with confidence due to low mappability). At the subfamily level, we observed higher expression of TEs on ZAL2^m^ than ZAL2 (*P* < 0.01, paired Mann–Whitney *U* test), which suggested potentially higher TE activity on ZAL2^m^. We also observed that TEs were more hypomethylated on ZAL2^m^ than ZAL2, and that this pattern was driven by ZAL2^m^ extremely hypomethylated DMCs (**Figure 3D**). Given that these effects were estimated at the TE subfamily level, more data are necessary to show a direct link between methylation of TEs and their insertion activity in the ZAL2^m^ chromosome.

### Potential regulatory consequences of ZAL2 and ZAL2^m^-specific DNA methylation

One of the best-known impacts of differential DNA methylation, when it occurs in promoters, is silencing of gene expression (2). Therefore, we first examined the expression levels of genes harboring allele-DMCs in their promoters. We found 325 genes with at least one allele-DMC in the promoter. For those genes, the divergence of gene expression was negatively correlated with the divergence of DNA methylation in the promoter (**Figure 4A**). This relationship was consistent with the aforementioned idea that promoter methylation dampens gene expression, although the degree of correlation was relatively weak (but significant). As we restricted our gene sets to those including more and more allele-DMCs in their promoters, the correlation coefficients increased (**Figures 4B-C**). These observations indicate that divergence in DNA methylation of promoters can explain some of the divergence in gene expression between ZAL2 and ZAL2^m^.

**Figure 4.**
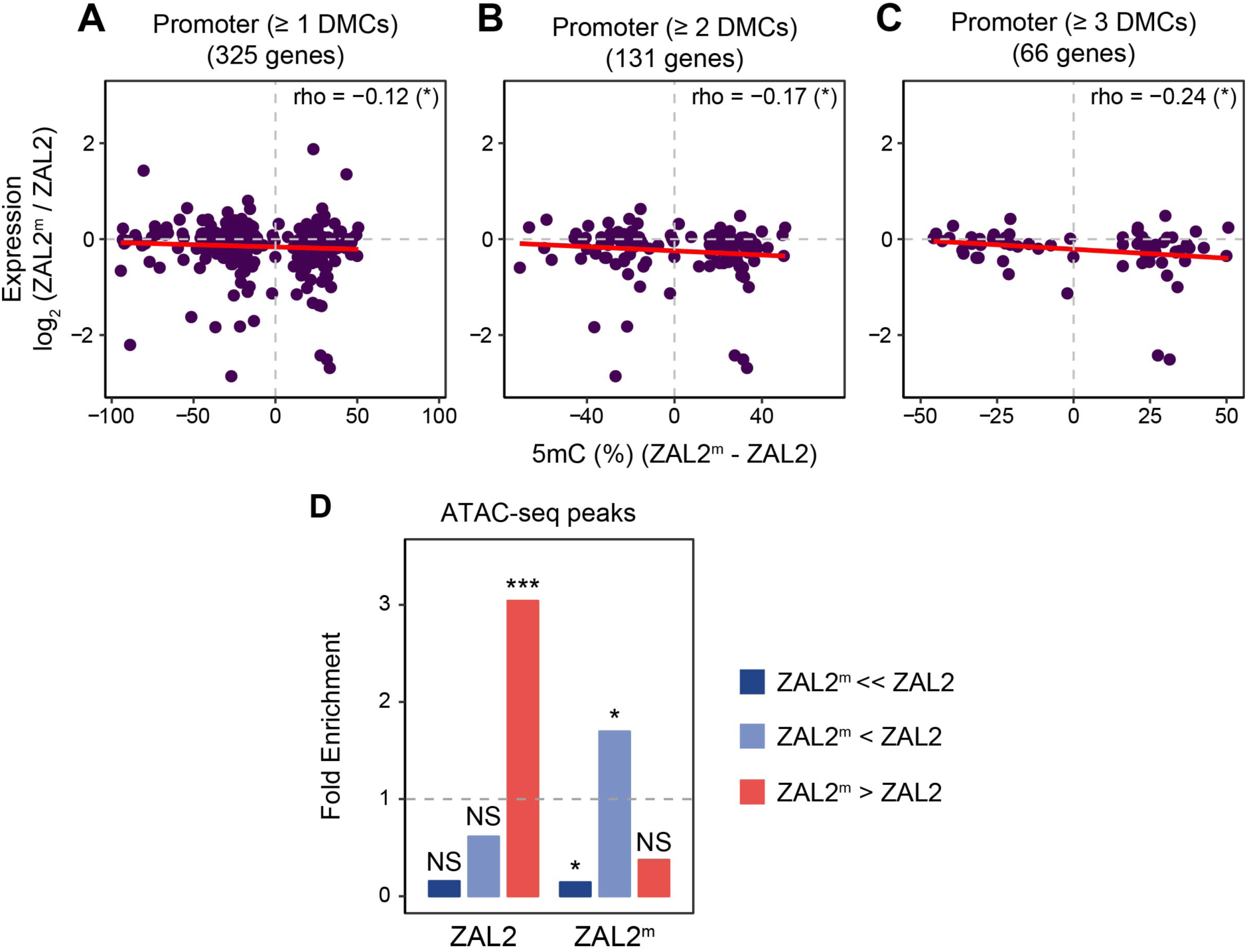
The potential role of allelic differences in DNA methylation in differential gene regulation. **(A-C)** Relationships between allelic differences in DNA methylation and allelic differences in gene expression for genes harboring more than 1, 2, and 3 allele-DMCs in their promoters. Allelic differences in DNA methylation across DMCs in a region were averaged. The strength and direction of association were measured by Spearman’s rank correlation coefficient, and the relationship was fit with a linear regression line (in red). **(D)** Fold enrichment of allele-DMCs occurring within ZAL2 or ZAL2^m^-specific ATAC-seq peaks. The dashed line corresponds to a fold enrichment of 1 (no enrichment or depletion relative to the background of all ZAL2/ZAL2^m^ CpGs). NS: not significant; \**P* < 0.05; \*\*\**P* < 0.001. Two proportion Z-test (also in Supplementary Table 2).

Recent studies have demonstrated that the regulatory impacts of differential DNA methylation extend far beyond promoters; differential DNA methylation can also cause differential expression of distant genes (60-63) and differential accessibility of long-range chromatin (64-67). We thus investigated the relationship between allele-DMCs and chromatin accessibility. We generated a map of accessible chromatin regions using ATAC-seq on DNA isolated from the hypothalamus of a white-throated sparrow of the WS morph (ZAL2/ZAL2^m^). We assigned open chromatin peaks to either ZAL2 or ZAL2^m^ (see Methods) and examined the overlap of each peak with allele-DMCs. In the absence of enrichment or depletion of allele-DMCs in these peaks, the number of ATAC-seq peaks that overlap allele-DMCs should be proportional simply to the number of CpGs, regardless of their allele-specific methylation status (Supplementary Table 2). In contrast to this prediction, we found both statistically significant enrichment and depletion of allele-DMCs within ATAC-seq peaks (**Figure 4D**, Supplementary Table 2). Only one ZAL2^m^ extremely hypomethylated allele-DMC was located within ATAC-seq peaks on each chromosome, which represents a significant (*P* = 0.037, proportion test) and marginally significant (*P* = 0.058) depletion of allele-DMCs from this category on ZAL2 and ZAL2^m^, respectively (**Figure 4D**). In contrast, other categories of allele-DMCs were enriched in allele-specific ATAC-seq peaks. Specifically, ZAL2^m^ hypomethylated allele-DMCs were enriched in ATAC-seq peaks specific to the ZAL2^m^ chromosome, but not in those specific to ZAL2 (**Figure 4D**). ZAL2^m^ hypermethylated allele-DMCs, on the other hand, were enriched in ZAL2 peaks but not in ZAL2^m^ peaks (**Figure 4D**). These observations suggest that differential DNA methylation between alleles correlates with their differential accessibility.

## DISCUSSION

Naturally occurring morphological and behavioral polymorphisms in white-throated sparrows offer a tremendous opportunity for studying the links between chromosomal differentiation and phenotypic traits (15-17). In this work, we present extensive epigenomic and transcriptomic data from this non-model organism, broadening our perspective on development and chromosomal evolution. We showed that developmental stages and plumage morphs are associated with distinct patterns of genome-wide DNA methylation in this species. The comparison between nestlings and adults revealed significant differences in DNA methylation that are widespread across the genome, except for the Z chromosome (**Figure 1C**). As previous studies of DNA methylation across development/aging have typically excluded sex chromosomes (e.g., (68,69), we have yet to understand why age-DMCs are underrepresented on the Z chromosome. Future studies that include sex chromosomes would reveal whether our observation is specific to white-throated sparrows or extends to other taxa.

Interestingly, age-DMCs were predominantly hypermethylated in adults (**Figure 2A**), consistent with the significantly higher expression of DNMTs in adults (**Figure 2B**). Previous studies have demonstrated widespread hypermethylation in the brains of humans and mice (12,70). As far as we are aware, however, changes in DNA methylation associated with aging have not been demonstrated outside of mammalian systems. Our observation of pronounced hypermethylation in adult brains, compared to nestling brains, in this avian species suggests that it may represent a shared molecular mechanism between mammals and birds. Previous studies in mammals have shown that DNA methylation regulates downstream pathways of neuronal and glial cellular differentiation (71-73), and that differential DNA methylation between neural cell types is critical for the differentiation of gene expression between them (41). Results of GO analysis and differential gene expression suggest that hypermethylation of promoter CpGs in adult brains might contribute to the down-regulation of early developmental genes (**Figure 2C**).

In contrast, morph differences in DNA methylation were nearly exclusive to the ZAL2/ZAL2^m^ chromosomes. We identified nearly 14,000 CpGs that were differentially methylated, at a relatively stringent cutoff of FDR-corrected *P* < 0.05. Utilizing these CpGs and outgroup data, we observed both hyper- and hypo-methylation of the non-recombining ZAL2^m^ chromosome as well as its counterpart, ZAL2, since their divergence. As DNA methylation varies strongly with underlying genetic variation in mammals and plants (74-77) some of the observed epigenetic divergence could have been due to divergence of linked positions. Most of the CpGs that were differentially methylated between the ZAL2 and ZAL2^m^ chromosomes were equally likely to be more methylated on either ZAL2 or ZAL2^m^. However, we discovered a group of CpGs that show extremely reduced DNA methylation on the ZAL2^m^ chromosome (referred to as ‘ZAL2^m^ extremely hypomethylated’ in the Results). This group accounted for a quarter of all allele-differentially methylated CpGs (**Figure 3**). Cross-species comparisons solidified that these CpGs underwent massive hypomethylation on the non-recombining ZAL2^m^ chromosome (**Figure 3B**).

Our study was designed to examine CpGs that are conserved on both the ZAL2 and ZAL2^m^ chromosomes, so that we could identify differentially methylated CpGs. It should be noted that CpGs that are specific to either chromosome might play important roles (17). For example, CpGs specific to humans are associated with cognitive traits and diseases (78), and are enriched in regions that are differentially DNA methylated between species (79). We intend to study the potential impacts of chromosome-specific CpGs in follow-up studies.

Experimental studies in human and mouse cell lines have demonstrated that recombination following double strand breaks can recruit DNA methyltransferases and increase DNA methylation (80-82). At the genome scale, methylation-associated SNPs and germline methylation levels are both positively correlated with inferred recombination rates in humans (83,84). Hypomethylation of the non-recombining chromosome in white-throated sparrows, ZAL2^m^, fits this broad observation, and supports a potential molecular link between recombination and DNA methylation. Interestingly, extreme hypomethylation of the ZAL2^m^ chromosome preferentially occurred in TEs (**Figure 3C**). ATAC-seq profiles of ZAL2^m^ extremely hypomethylated CpGs indicate that they tend to occur outside of accessible chromatin (**Figure 4D**). Hypomethylation is known to activate TEs (85), further increasing TE insertion (86,87). We showed that at the subfamily level, TEs on ZAL2^m^ exhibit higher expression than those on ZAL2, which is consistent with the effects of hypomethylation on TE activity (**Figure 3D**). Given that an increase in TE insertion is hypothesized to be one of the first genomic changes during the evolution of non-recombining chromosomes in *Drosophila* (88), a similar mechanism may be operating in the ZAL2/ZAL2^m^ system, potentially fueled by the extreme hypomethylation. Additional data on TE transcription and a better-annotated reference genome in this species will be necessary to investigate the relationship between DNA methylation and TE activity on the ZAL2^m^ chromosome.

Integrating our gene expression data and chromatin accessibility data, we present results consistent with regulatory roles of allele-specific DNA methylation. First, when allele-DMCs were present in the promoter, the degree of differential methylation of those promoters was correlated with the degree of differential expression of the genes (**Figure 4A-C**). Second, the landscape of open chromatin on the ZAL2 and ZAL2^m^ chromosomes in a WS bird suggested significant associations between allele-specific hypomethylation and allele-specific open chromatin peaks (**Figure 4D**). The comparison between ATAC-seq peaks and DNA methylation should be taken with caution because of a limitation in our data; the tissue sample used for ATAC-seq was from a non-breeding (winter) male while the adult WGBS data were from breeding (summer) females. ATAC-seq and WGBS data from the same birds are currently lacking. A recent study of 66 ATAC-seq maps from 20 different tissues of male and female mice (89) demonstrated that the majority of accessible regions between tissues overlapped and that the correlation between male and female tissues was extremely high. For example, in samples of cerebellum in mice, the correlation in accessible regions between males and females was 0.96 in (89)). In the present study, the associations between DNA methylation and chromatin accessibility are consistent with those observed in model organisms (64-66) and suggest that changes in allele-specific DNA methylation may correlate with the chromatin landscape. Together, these observations indicate widespread functional impacts of differential DNA methylation in the genome of this interesting species.

In conclusion, our comprehensive epigenetic study in white-throated sparrows has revealed significant effects of age and plumage morph on DNA methylation landscapes. We show that effects of age on DNA methylation are pervasive and likely affect regulation of developmental genes. In contrast, morph differences in DNA methylation are mostly enriched on ZAL2/ZAL2^m^, and involve both hyper- and hypomethylation of the recombination-suppressed ZAL2^m^ as well as its counterpart, ZAL2. On the basis of a comparison with an outgroup, we also discovered a large number of CpGs for which DNA methylation has been dramatically reduced specifically on ZAL2^m^ chromosome. We propose that these different varieties of allelic DNA methylation divergence have led to specific functional consequences. Together, our results not only provide a novel data set from a wild avian species, but also raise several hypotheses on which we hope future studies will build to further illuminate the connection between genotype and phenotype and pathways of chromosome evolution.

## ACKNOWLEDGEMENTS

This study was funded by an NIH grant (R01MH082833) to DLM and an NSF grant (IOS-1656247) to DLM and SVY, and by NIH (R01MH103517) and NSF (MCB-1615664) grants to SVY. We thank Ben Long for comments on the manuscript.

## Notes

### Competing Interest Statement

The authors have declared no competing interest.

